# Incorporating special interests to investigate the language system in autism: A feasibility pilot fMRI study

**DOI:** 10.1101/2025.04.04.647117

**Authors:** Halie A. Olson, Kristina T. Johnson, Shruti Nishith, Anila M. D’Mello

## Abstract

Most autistic individuals have sustained, focused interests in particular topics or activities. In some cases, these special interests have been shown to motivate communicative behaviors, a domain in which many autistic individuals experience challenges. We conducted a pilot study with 15 autistic children (ages 8.18 – 13.27 years, mean(SD)= 11.17(1.62), 3 female/11 male/1 nonbinary), comparing brain responses elicited by short narratives tailored to individuals’ special interests to responses elicited by generic, non-tailored narratives. Using functional magnetic resonance imaging (fMRI), we found that autistic children did not show typical language responses to generic narratives. However, they did show heightened responses to the narratives that incorporated their special interests relative to the generic narratives in language regions and in regions associated with reward and self-reference. Brain responses for personalized narratives were also more consistent across children than responses for the generic narratives. These results suggest that personalizing stimuli by incorporating special interests might be a promising approach for neuroimaging in autistic participants.

**LAY SUMMARY:** In a pilot sample of autistic children, we found that listening to short narratives tailored to each child’s special interest elicited higher responses in the brain than listening to non-personalized narratives. Brain responses across children were also more similar for the special interest narratives. Thus, personalizing stimuli to special interests may be a promising approach for neuroimaging studies of autism.

## INTRODUCTION

Difficulties with language and communication are a consistent characteristic of autism, enduring across the lifespan and affecting an individual’s well-being and global functioning (Howlin, 2003; Miranda et al., 2023). However, the brain basis of these language difficulties remains elusive. One reason is that traditional language neuroimaging paradigms are not optimized to account for the unique challenges faced by many autistic individuals. Most fMRI language paradigms embed cognitive demands and stimuli that were originally designed for and normed on neurotypical participants. Further, autism is a highly heterogeneous condition, making it difficult to devise paradigms that are equally accessible, engaging, and relevant across individuals (Jeste & Geschwind, 2014; Tager-Flusberg, 2004).

In our prior research in neurotypical children, we investigated the efficacy of personalizing neuroimaging paradigms to study language (Olson, Johnson, et al., 2024). To do this, we identified a topic of high interest for each child and recorded narratives based on this interest. When comparing activation for these personalized interest narratives to generic narratives that were the same across individuals, we found stronger activation and more inter-subject consistency in language responses for personalized narratives. These findings not only demonstrate a modulatory effect of interest on the language system, but also show that using interests to personalize neuroimaging stimuli is a feasible, effective, and promising method for studying the brain basis of language.

This approach is particularly well-suited to studying language in autism. A core diagnostic feature of autism is the presence of highly focused “special interests” (i.e., restricted/circumscribed interests, affinities; American Psychiatric Association, 2013). Special interests are highly prevalent and persistent across the lifespan in autism (Grove et al., 2016; Klin et al., 2007; Spackman et al., 2023). Incorporating special interests into therapeutic settings has been associated with better social communication in autism (Arunachalam et al., 2024; Baker et al., 1998; Boyd et al., 2007; Charlop-Christy & Haymes, 1998; Harrop et al., 2019; Lizon et al., 2023), and fMRI studies of autism that use special interests as personalized stimuli find increased activation in regions sometimes found to be underactive in autism (Antezana et al., 2022; Cascio et al., 2014; Foss-Feig et al., 2016; Kohls et al., 2018).

Taken together, incorporating special interests in an fMRI study design may provide a window of opportunity to examine language in a way that reduces demands on participants and is sensitive to the inherent heterogeneity of autism. Here, we test this approach in a pilot study of n=15 autistic children.

## MATERIALS AND METHODS

### Participants

Data were analyzed from 15 autistic children (3 female/11 male/1 nonbinary; **Table 1**). All children were native English speakers, had normal hearing, normal or corrected-to-normal vision, no contraindications for MRI, and a qualifying special interest (i.e., must (1) engage with the interest for an average of an hour per day, or would engage without screen time restrictions, (2) have had the same interest for at least the last two weeks, and (3) have an interest that could be represented in video format). Verbal and nonverbal cognitive reasoning were assessed via the Kaufman Brief Intelligence Test (KBIT-2; Kaufman & Kaufman, 2004). Caregivers completed questionnaires about their child including demographic and developmental histories (e.g., language onset) and the Social Responsiveness Scale (SRS; Constantino & Gruber, 2005). In n=9 autistic children scanned prior to the COVID-19 pandemic, diagnosis was confirmed via the Autism Diagnostic Observation Schedule (ADOS) administered by a research-reliable clinician (Lord et al., 2000); remaining children had received a formal diagnosis from a qualified provider and had SRS Total scores in the moderate to severe range, which are strongly associated with a diagnosis of autism. N=13 additional autistic children were recruited but did not contribute usable data (n=5 refused to enter the scanner or were unable to stay in the scanner past the initial T1, n=8 had excessive motion).

**Table 1:** Demographics and interests. Group means (and standard deviations) for several demographic and cognitive measures. KBIT = Kaufmann Brief Intelligence Test, SRS = Social Responsiveness Scale. One participant did not complete the KBIT and is not included in the table above. Another participant did not complete the KBIT at the scanning visit, so KBIT scores come from a prior visit. Language onset measures are based on caregiver report.

| Demographic information |  |
| --- | --- |
| Age (years) | 11.17(1.62), range = 8.18 – 13.27 |
| KBIT Matrices (standard score) | 114.93(7.92), range = 104.00 – 133.00 |
| KBIT Verbal Composite (standard score) | 107.57(12.70), range = 77.00 – 119.00 |
| SRS Total (T score) | 77.75(15.2), range = 48 – 101 |
| Age of first word (months) | 14.00(8.32), range = 8 – 42 |
| Age of first sentence (months) | 23.00(7.78), range = 12 – 42 |
| Age for understanding command (months) | 14.93(9.64), range = 3 - 36 |
| Interests | Soccer<br>Tennis<br>Computers<br>Fortnite<br>Minecraft<br>Pokémon<br>Among Us video game<br>Lego Ninjago (n=2)<br>Cartoon voices<br>Dragons from How to Train Your Dragon<br>Fighting insects<br>Puppies<br>Hurricanes/Extreme weather<br>Trains (local commuter line) |

### Experimental Protocol

Participants completed behavioral testing and MRI scanning across 1-2 visits. The MRI session included an anatomical scan, a functional task in which participants watched their special interest videos and nature videos (not discussed in this paper), the special interest language task, and optional additional scans beyond the scope of the current study. See **Supplement** for MRI acquisition and modeling details.

### Special Interest Language fMRI Task

Stimulus creation and task details have been previously reported in Olson, Johnson, et al. (2024) and are briefly summarized here. Narratives were written based on video clips of nature scenes (NEUTRAL condition) and special interest videos provided by families (INTEREST condition) that children watched prior to completing the language task. Special interest narratives were highly tailored to each child based on provided materials (e.g., using specific terminology, focusing on timestamps identified as favorite parts of the videos). All narratives were recorded by the same female experimenter, and were approximately matched between participants by avoiding personal pronouns, using simple vocabulary (allowing for interest-specific terms), and using short sentences. See OSF (https://osf.io/dh3wq/) for detailed information on narratives including transcripts.

During the fMRI task, children listened to 21 narratives (16 seconds each) while viewing a fixation cross, with a 5-second fixation rest block between each narrative. In the INTEREST condition, participants listened to the personalized special interest narratives. In the NEUTRAL condition, participants listened to non-personalized narratives describing nature. In the BACKWARDS condition, participants listened to incomprehensible reversed versions of neutral narratives to account for low-level auditory features of the narratives. A low-demand attention check (press button when panda appeared) occurred after each narrative. Condition order was fixed across participants in an [ABCABC…] pattern, alternating INTEREST, NEUTRAL, then BACKWARDS (**Figure 1**).

**Figure 1:**
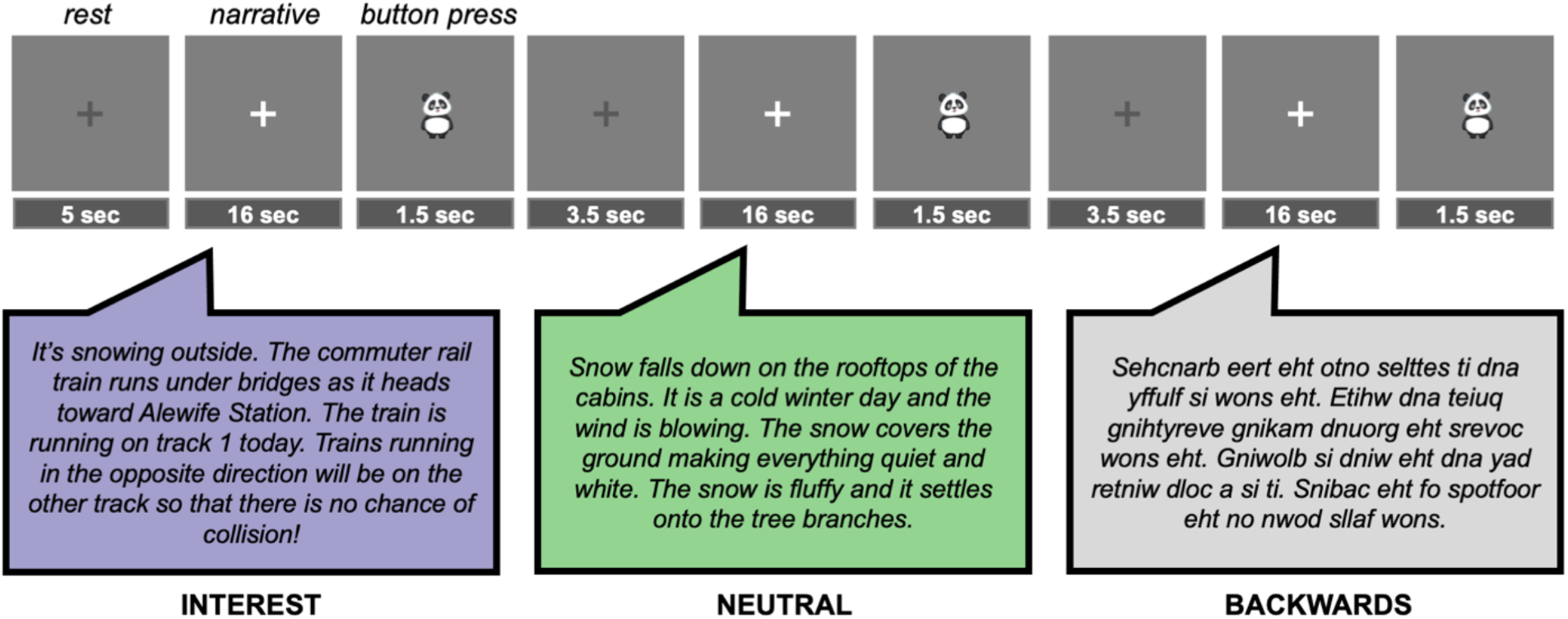
Task design and example narratives. Figure from Olson, Johnson, et al., 2024.

### fMRI Analyses

Parameter estimates for each condition were extracted from *a priori* regions of interest (ROIs) important for language processing (Lipkin et al., 2022): left IFGorb, left IFG, left MFG, left AntTemp, left PostTemp, left AngG, right cerebellum lobule VI, and right cerebellum Crus I/II. Linear mixed-effects models were run in R’s `lme4` package. To determine if there was an effect of condition (INTEREST, NEUTRAL, BACKWARDS), we used:

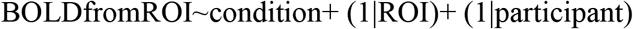

with participant and ROI as random factors to account for repeated measures. Post-hoc tests were then conducted using the `lsmeans` package.

We also conducted whole-brain analyses for contrasts of interest (NEUTRAL, INTEREST, INTEREST > NEUTRAL) using one-sample t-tests. Results were thresholded at an uncorrected voxel p < 0.001, with a cluster corrected FWE < 0.05.

### Jaccard Index

Inter-individual similarity in the spatial distribution of the whole brain responses for INTEREST and NEUTRAL conditions was assessed using the Jaccard Index (JI) (Jaccard, 1908). JI is a measure of the proportion of voxels in common between two conditions with respect to the total of activation voxels identified by either condition. In each condition, each participant’s first-level activation map was thresholded at t>2.3 and binarized. JI was calculated across the whole brain using the formula: Map_i_∩Map_j_/Map_i_∪Map_j_ for all possible pairs of maps *i* and *j*.

## RESULTS

### Response to special interests and neutral language in language regions

The primary goal of the study was to determine the utility of using special interests to study language processing in the brain in autism. We first examined the magnitude of the response to INTEREST, NEUTRAL, and BACKWARDS conditions within canonical language regions (**Figure 2A**). Neurotypical children typically show a robust difference in brain activation between language and a lower-level control condition (e.g., backwards speech) in these regions (e.g., Ozernov-Palchik, O’Brien, et al., 2024), including in our prior work on this task in a sample of age-matched neurotypical children (Olson, Johnson, et al., 2024). Here, we found that activation in language regions in autistic children differed by condition (F(2,336) = 47.71, p < 0.001). Notably, participants showed no significant difference in activation between the NEUTRAL and BACKWARDS conditions in language regions (NEUTRAL>BACKWARDS: t(336) = 1.17, p = 0.47) consistent with prior literature showing atypical language processing in autism. However, special interest narratives elicited higher activation than both neutral and backwards narratives in language ROIs (INTEREST>NEUTRAL: t(336) = 7.81, p < 0.001; INTEREST>BACKWARDS: t(336) = 8.98, p < 0.001; **Figure 2A**).

**Figure 2:**
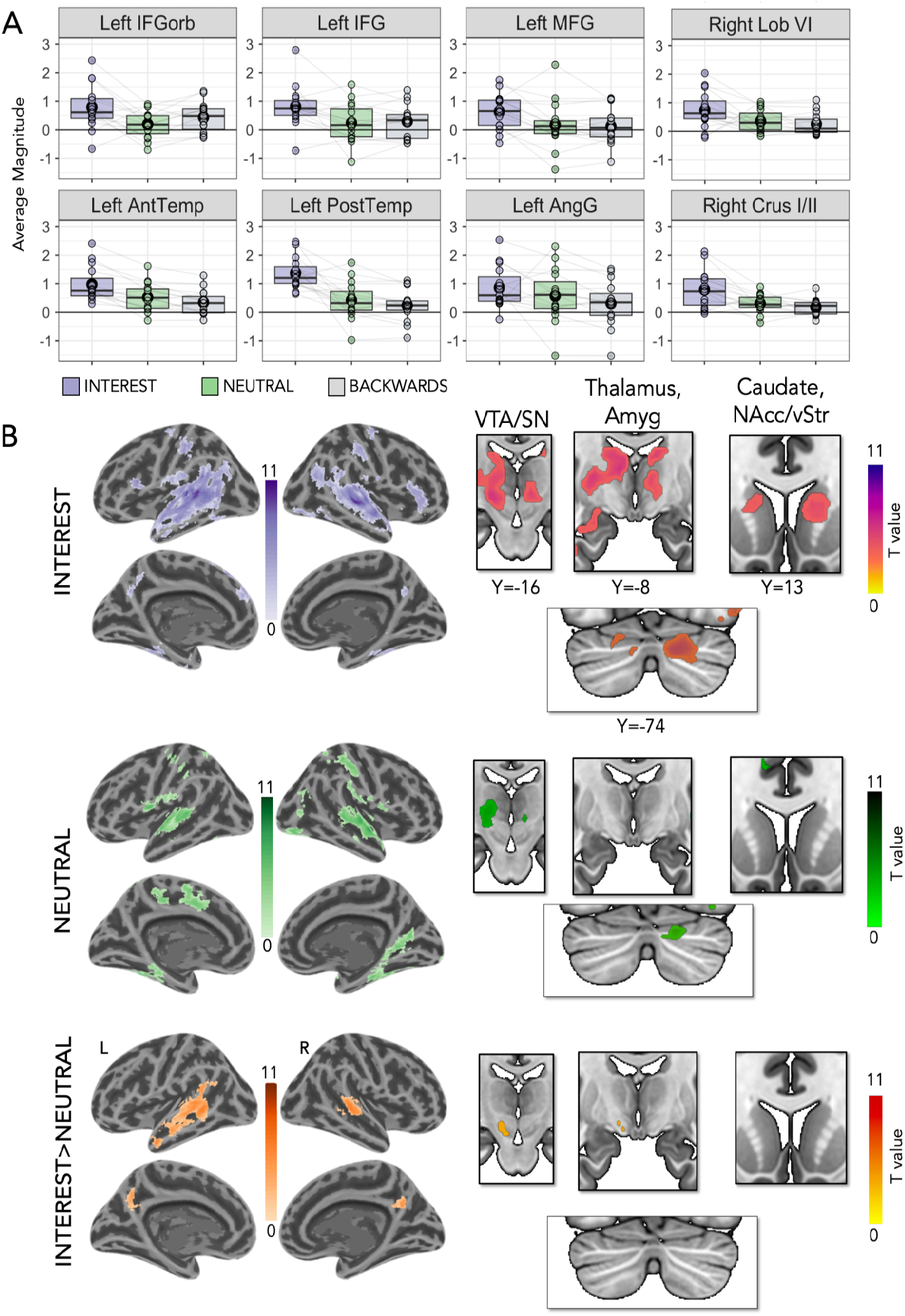
Special interest language increases brain activation in autistic children. (A) Boxplots show average magnitude per condition in each a priori language region of interest. Circles represent individual means, and lines connect individual participants across conditions. (B) Whole-brain group activation maps for the INTEREST>NEUTRAL, INTEREST>Baseline, and NEUTRAL>Baseline (n=15). Threshold: p<0.001, FWE cluster corrected at p<0.05.

We also expected that special interest narratives would activate regions beyond canonical language regions, based on prior literature (Cascio et al., 2014; Foss-Feig et al., 2016; Kohls et al., 2018) and our prior work (Olson, Johnson, et al., 2024). We thus additionally conducted a group-level whole-brain analysis. Whole-brain analyses identified higher responses to INTEREST than NEUTRAL conditions in bilateral temporal regions and regions involved in reward (e.g., ventral tegmental area/VTA) and self-referential processing (e.g., precuneus) (**Figure 2B**). These results were observable both at the group level (**Figure 2B**) and in individual participants (**Figure 3A;** for individual responses in all participants, see **Supplement**).

**Figure 3:**
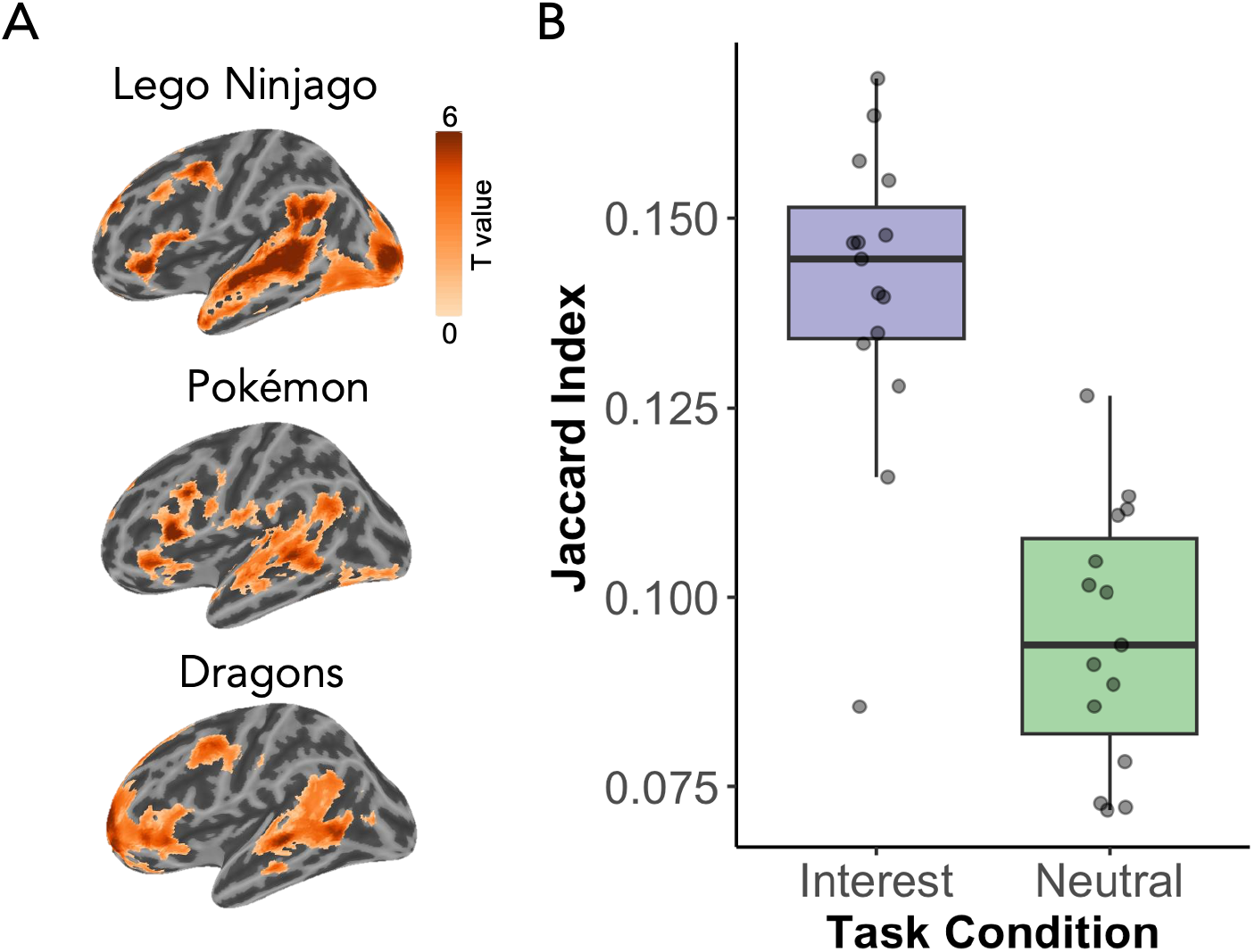
Greater consistency in brain responses across individual autistic children for special interest narratives. (A) Individual whole-brain responses from n=3 representative participants to INTEREST>NEUTRAL language visualized at p<0.01, FWE cluster p<0.05. Participants activate consistent regions despite idiosyncratic language stimuli. (B) Jaccard indices quantifying intersubject consistency (for each pair of participants) in whole brain activation patterns for the INTEREST (purple) and NEUTRAL (green) conditions. Each dot represents the mean JI for each participant.

### Consistency in language responses for special interest versus neutral language stimuli

One potential downside of using idiosyncratic stimuli across participants is the possibility of greater between-participant variation in the spatial distribution of the neural response. Our prior research in neurotypical children found *greater* consistency in neural responses when children listened to highly diverse personalized narratives than neutral narratives (Olson, Johnson, et al., 2024). To investigate this in autistic children, we computed the Jaccard Index - a measure of spatial consistency between participants. Autistic children showed more consistent activation patterns for INTEREST over NEUTRAL narratives (t(14)=9.06, p<0.0001), despite the fact that special interest narratives differed across participants when neutral narratives did not (**Figure 3B**).

## DISCUSSION

In a pilot sample of autistic children, we found that tailoring fMRI language stimuli to each child’s special interest had a positive effect on activation in the language system. Consistent with prior studies finding atypical language activation in autism (Eyler et al., 2012; Herringshaw et al., 2016; Kleinhans et al., 2008; Lindell & Hudry, 2013; Lombardo et al., 2015; Mody & Belliveau, 2013), our participants did not show the expected difference between the generic versus backwards narratives, even though neurotypical children do show this difference (see Olson, Johnson, et al., 2024 for typical generic > neutral results on this task). This result makes it seem as if language regions were not engaged strongly by language comprehension in autism. However, when comparing special interest narratives to generic or backwards narratives, these same language regions were highly activated, suggesting that the *content* of language has a powerful effect on activation within the language system in autism.

Why might this be the case? Autistic individuals engage with special interests often and deeply and find them highly motivating (Grove et al., 2016, 2018). Thus, increased brain activation for special interests (in language regions and in salience, reward, and self-reference regions) may be the result of higher attention to or increased familiarity with stimuli (see Olson, Johnson, et al., 2024 for discussion). In a handful of studies in which experimental stimuli were tailored to autistic participants (e.g., mother’s face, special interests), researchers found increased brain activation in regions commonly underactive in autism, such as face processing and reward regions (Antezana et al., 2022; Cascio et al., 2014; Foss-Feig et al., 2016; Kohls et al., 2018; Pierce & Redcay, 2008). Critically, potentiating effects of interests on the brain are not autism-specific: interests also potentiate language activation in neurotypical children (Olson, Johnson, et al., 2024). However, these findings are particularly notable in autism because they might provide a way to probe systems that are typically underactive in autism (e.g., language/reward), and ultimately influence the interpretation of their capacity or function.

Notably, we also found that idiosyncratic special interest narratives resulted in *more* consistent brain activation patterns than generic narratives. This may be because participants varied in their level of attention to or engagement with generic narratives (resulting in less consistent activation across autistic individuals), whereas special interest narratives were chosen to be maximally engaging to each participant.

Incorporating special interests into research is not only effective but also feasible. Up to almost 90% of autistic children (Klin et al., 2007) have special interests (which persist even into adulthood), making it easy to identify appropriate stimuli for each participant. Furthermore, the advent of new technologies (such as generative artificial intelligence models and human-like text-to-speech, which can be integrated into stimuli development pipelines; e.g., Santi et al., 2025) make it possible to scale up personalized designs such as this one to larger studies and even clinical contexts.

Finally, our results add to a growing consensus that autism research should be sensitive to the diverse ways that autistic individuals experience the world (Dwyer, 2022; Pellicano & den Houting, 2022). Incorporating special interests is consistent with a ‘strengths-based’ design: it explicitly incorporates participants’ unique knowledge and reduces barriers to successful participation in research. A compelling possibility to be explored in future research is that study designs centered on neurotypical norms might not reflect the true capacity of autistic individuals. This is particularly relevant to the clinical domain. Clinical studies (Arunachalam et al., 2024; Baker et al., 1998; Boyd et al., 2007; Charlop-Christy & Haymes, 1998; Harrop et al., 2019; Lizon et al., 2023) and anecdotal reports (Suskind, 2014) describe positive effects of incorporating special interests on social communication. While in our current study we cannot say whether higher activation is associated with “better” language processing, our results do suggest that special interests have a powerful modulatory effect on the brain.

In conclusion, here we assess a personalized neuroimaging approach incorporating special interests in a pilot sample of 15 autistic children, adding to the emerging evidence that special interests provide a window of opportunity to investigate brain function in autism. Future work should aim to expand such an approach to larger sample sizes and other cognitive domains and age groups.

## ACKNOWLEDGEMENTS

We thank Isabelle Frosch and Cindy Li for assistance with recruiting, coordinating, and testing. We also thank Hannah Grotzinger for task programming assistance, Caitlin Malloy for ADOS support, and our undergraduate and high school research assistants who assisted with various aspects of the project, including Jimmy Chen, Nicole Dundas, Insha Merchant, Rucha Kelkar, Alana Kalehua, and Hillary Jean-Gilles. We are grateful to Atsushi Takahashi and Steve Shannon from the Athinoula A. Martinos Imaging Center at MIT. We thank the participants and families for making this research possible. Finally, we thank Ron Suskind, whose experience with his son Owen inspired this research.

