## Supplement for "Incorporating special interests to investigate the language system in autism: A feasibility pilot fMRI study"

#### Supplemental Methods

**Preregistration**. The main hypotheses for the current paper were included as part of a broader preregistration in 2018 for a study investigating the neural correlates of personal interest neurotypical and autistic children: https://osf.io/nr3gk. We deviated from the preregistration by not using subject-specific functional ROIs (NEUTRAL>BACKWARDS), as this would have precluded a comparison between our conditions of interest (INTEREST vs. NEUTRAL). We did not test hypotheses related to associations between neural and behavioral measures, due to smaller than anticipated sample sizes as a result of the COVID-19 pandemic and related data acquisition and personnel limitations.

**Stimuli Features.** We tested several sub-lexical and lexical features of narratives to ensure matching across INTEREST and NEUTRAL conditions as possible. We conducted non-parametric pairwise comparisons of several extracted features using Wilcoxon rank-sum tests to identify specific differences between each individual participant's narratives and the neutral narratives (see **Supplemental Figure 1**). After applying a Bonferroni correction for multiple comparisons (adjusted $p<0.0033$ for 15 comparisons), no significant differences were found for the number of sentences, emotional valence, adverb usage, verb usage, and adjective usage across narratives. Some subjects showed significant differences between their personalized narratives and the set of neutral narratives in the number of words, the number of syllables per sentence, average word frequency, and noun and proper noun usage. Because the NEUTRAL condition contained no proper nouns, we also combined nouns and proper nouns in the personalized narratives and then found no significant differences across all nouns. Word frequency and proper noun usage are expected to differ given that special interest vocabulary may be more specific (see Olson, Johnson, et al., 2024 for discussion).

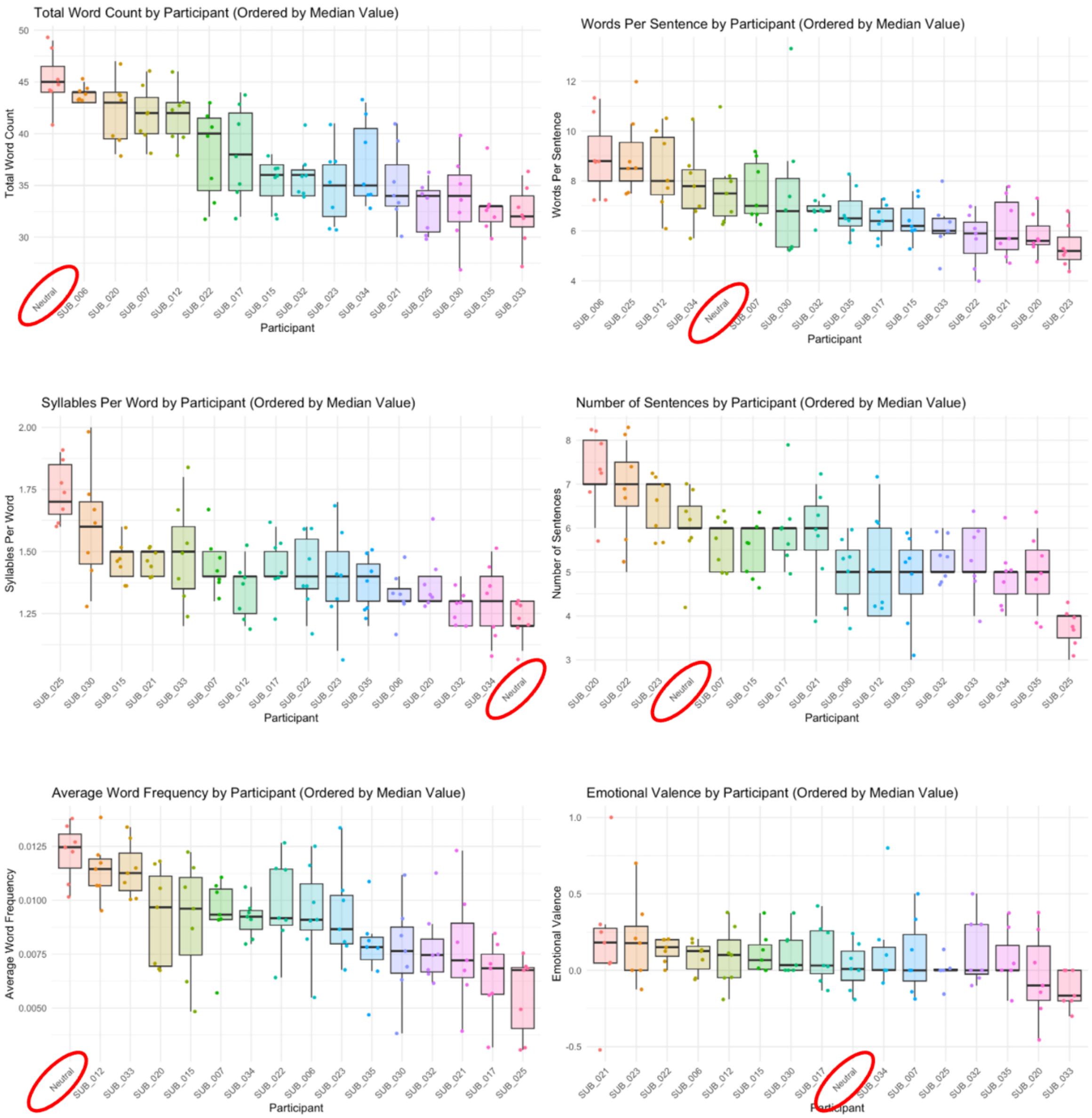
Total Word Count by Participant (Ordered by Median Value)
Total Word Count
50
45
40
35
30
Neutral
SUB_006
SUB_020
SUB_007
SUB_012
SUB_022
SUB_017
SUB_015
SUB_032
SUB_023
SUB_034
SUB_021
SUB_025
SUB_030
SUB_035
SUB_033
Participant
Words Per Sentence by Participant (Ordered by Median Value)
Words Per Sentence
12
10
8
6
4
SUB_006
SUB_025
SUB_012
SUB_034
Neutral
SUB_007
SUB_030
SUB_032
SUB_035
SUB_017
SUB_015
SUB_033
SUB_022
SUB_021
SUB_020
SUB_023
Participant
Syllables Per Word by Participant (Ordered by Median Value)
Syllables Per Word
2.00
1.75
1.50
1.25
SUB_025
SUB_030
SUB_015
SUB_021
SUB_033
SUB_007
SUB_012
SUB_017
SUB_022
SUB_023
SUB_035
SUB_006
SUB_020
SUB_032
SUB_034
Neutral
Participant
Number of Sentences by Participant (Ordered by Median Value)
Number of Sentences
8
7
6
5
4
3
SUB_020
SUB_022
SUB_023
Neutral
SUB_007
SUB_015
SUB_017
SUB_021
SUB_006
SUB_012
SUB_030
SUB_032
SUB_033
SUB_034
SUB_035
SUB_025
Participant
Average Word Frequency by Participant (Ordered by Median Value)
Average Word Frequency
0.0125
0.0100
0.0075
0.0050
Neutral
SUB_012
SUB_033
SUB_020
SUB_015
SUB_007
SUB_034
SUB_022
SUB_006
SUB_023
SUB_035
SUB_030
SUB_032
SUB_021
SUB_017
SUB_025
Participant
Emotional Valence by Participant (Ordered by Median Value)
Emotional Valence
1.0
0.5
0.0
-0.5
SUB_021
SUB_023
SUB_022
SUB_006
SUB_012
SUB_015
SUB_030
SUB_017
Neutral
SUB_034
SUB_007
SUB_025
SUB_032
SUB_035
SUB_020
SUB_033
Participant

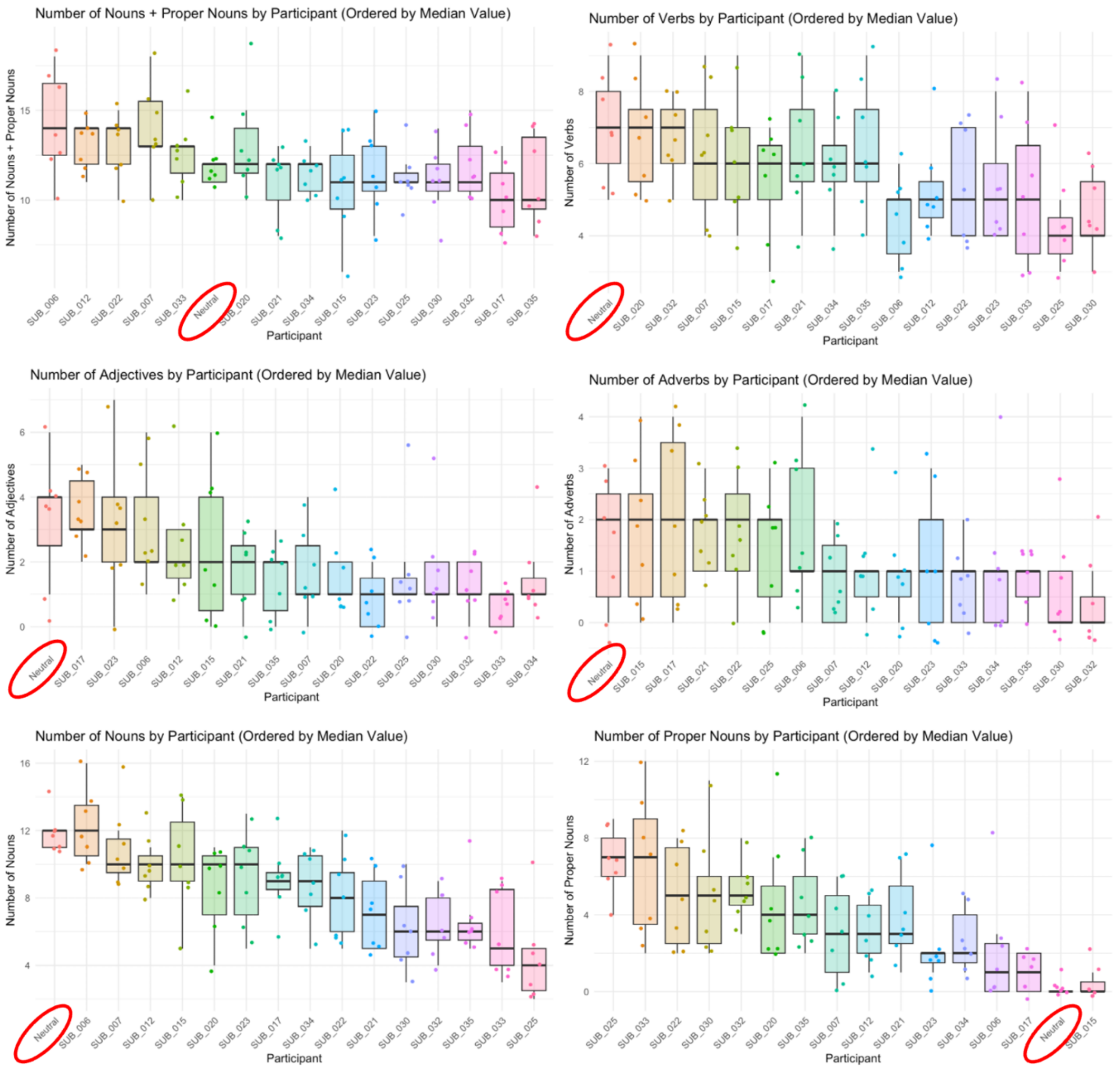


***Supplementary Figure 1: Linguistic and paralinguistic features for individual participants' stimuli.*** *Box plots show several features of the narratives. Individual participants are shown in different colors and each narrative is represented by a dot. Plots are ordered by median values. The neutral condition is circled in red for reference.*

**MRI Acquisition**. Data were acquired from a 3-Tesla Siemens Prisma scanner located at the Athinoula A. Martinos Imaging Center at the McGovern Institute at MIT, using a 32-channel head coil. T1-weighted structural images were acquired in 196 interleaved slices with 1.0mm isotropic voxels (MPRAGE; TA=5:08; TR=2530.0ms; FOV=256mm; GRAPPA parallel imaging, acceleration factor of 3). Functional data were acquired with a gradient-echo EPI sequence sensitive to Blood Oxygenation Level Dependent (BOLD) contrast in 3.0mm isotropic voxels in 40 near-axial slices covering the whole brain (EPI factor=70; TR=2500ms; TE=30ms; flip angle=90 degrees; FOV=210mm; TA=7:47).

**Preprocessing and Statistical Modeling**. fMRI data were preprocessed in fMRIPrep v1.1.1 (Esteban et al., 2019) and smoothed at 6mm FWHM. First level modeling was performed using SPM12 with individual regressors for INTEREST, NEUTRAL, BACKWARDS, and button press. TRs were marked as outliers if they had greater than 1mm of framewise displacement. We excluded participants with > 20% outlier volumes.

### Supplemental Results

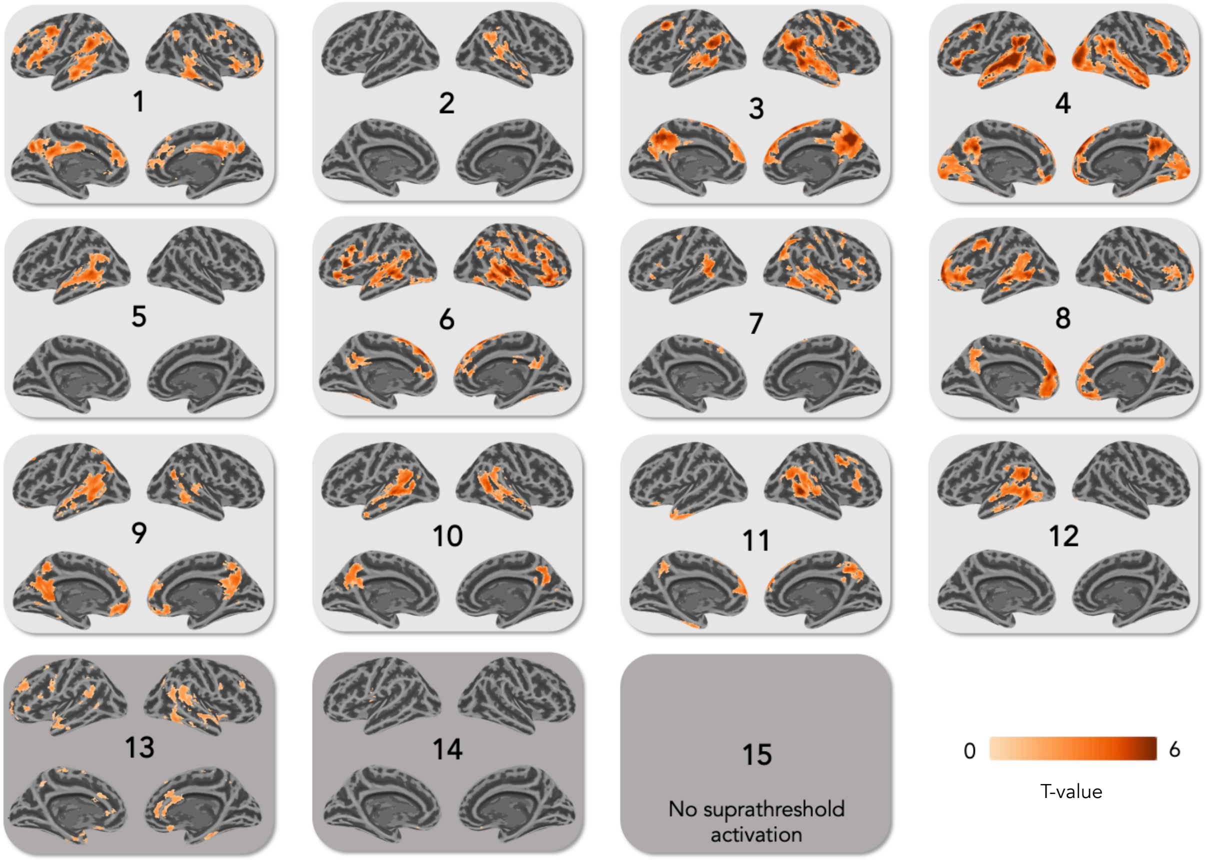


***Supplementary Figure 2: Higher responses to special interest narratives in individual autistic participants***. *Individual whole-brain responses from 15 autistic participants to INTEREST>NEUTRAL language. N=12 participants visualized at p<0.01, FWE cluster p<0.05. N=2 participants did not show activation at this threshold or activation was not well visualized in the surface rendering and are thus visualized at p<0.05 uncorrected, and n=1 participant did not show activation at the uncorrected threshold (p<0.05).*
